# Multimodal electrical impedance tomography and electroencephalography imaging: Does higher skull conductivity resolution in EIT imaging improve accuracy of EEG source localization?

**DOI:** 10.1101/2024.08.05.606582

**Authors:** Ville Rimpiläinen, Alexandra Koulouri

## Abstract

**Objective:** Unknown conductivities of the head tissues, particularly the skull, is a major factor of uncertainty in electroencephalography (EEG) source imaging. Here, we develop a personalized skull conductivity framework aiming to improve the head models used in the EEG source imaging and to reduce localization errors.

**Methods:** We employ Electrical Impedance Tomography (EIT) and convex optimization to produce high resolution skull conductivity maps that are subsequently embedded in the EEG modeling.

**Results:** First, we demonstrate through simulations that locally varying conductivity values of the skull can be estimated from EIT measurements. Second, we show how the choice of the skull conductivity resolution of the EIT imaging affects the EEG source reconstructions.

**Conclusions:** EIT estimated conductivities can signicantly improve the source reconstructions, particularly in cortical areas under bones that exhibit high conductivity variations.

**Significance:** This work acts as a steppingstone in defining a protocol for the preparation of patient-specic head conductivity models that are essential for accurate examination and systematic monitoring of the brain activity via EEG.

## 1 Introduction

The determination of the electric conductivity values of the different tissues of the human head is an important and challenging problem that is often encountered in electroencephalography (EEG) source imaging in which active parts of the brain are determined based on scalp potentials. The EEG source imaging results depend strongly on the conductivity modelling of the tissues, particularly the highly insulating skull [42, 9, 24, 29, 37].

A recent review showed that there is a wide range of conductivities reported in literature for the whole (bulk) skull: the values ranged from 0.00182 S/m to 1.718 S/m with a weighted mean value 0.016 S/m [27, 28]. In another study, a significant inter-subject variability in skull conductivity was observed with reported mean value and standard deviation of 0.00844 ± 0.00484 S/m [3]. It seems that age is a significant factor to explain the inter-subject variability [17, 27, 3]. In addition to inter-subject variability, there is also variability in the whole skull conductivity within the same subject. For example, in [23] it was reported that the electric conductivity of the temporal bones can be much lower (lowest measured value 0.0047 S/m) than elsewhere, e.g. in the occipital bone (highest measured value 0.0735 S/m). Here, the values depended on whether the measured sample contained sutures and cancellous bone, and in addition also a correlation to the thickness of the sample was reported [23]. Therefore, a tool that can solve the skull conductivities individually is needed.

Currently, there are only few techniques to determine (or calibrate) head tissue conductivities *in vivo*. These techniques usually utilize either well defined somatosensory evoked potentials / fields (SEP/SEF) in combination with EEG [25, 34], EEG/MEG [12, 19, 47, 5, 4, 3] or electrical impedance tomography (EIT) [13, 14, 8, 10]. The main limitation of using EEG or EEG/MEG with SEP/SEF data is that these measurements can only be used for a single parameter (bulk skull conductivity) calibration, and for example scalp conductivity has to be known. In addition, the SEP/SEF calibrated skull conductivity might not be optimal for source activity that locates in another brain region [34].

Electrical impedance tomography (EIT) is an imaging modality that can be used to solve internal conductivity profiles with the help of boundary electrodes [18]. Most EIT studies, however, have been rather limited in the sense that only very few, 4 or less, (bulk) tissue conductivity parametres have been solved [13, 14, 8, 10]^1^. This effectively has neglected any possibility to find local variations within any of the tissue compartments. In a recent study, it was showed that it was possible to image locally varying skull conductivity values with roughly 3 - 4 cm resolution [11]. One of the main motivations for using EIT to image locally varying skull conductivity has been that these conductivities could improve EEG source localization. However, to the best of our knowledge, there are no systematic studies in which this benefit is quantified. Therefore, two questions remain: 1) how much EIT skull conductivity imaging can improve EEG source localization; 2) with how fine resolution the skull conductivity should be imaged in order to be beneficial for EEG source imaging.

In our preliminary study [38], we demonstrated that different conductivities in a five-piece skull can be reconstructed with the help of a box constraint. For the present study, we constructed a numerical head phantom in which the skull conductivity values are smoothly varying based on measured conductivities of 20 skull samples drilled from a postmortem human skull [23].

The contributions of our present study are the following:

- We propose an EIT conductivity imaging algorithm that employs convex optimization to reconstruct (low and) high-resolution skull conductivity maps of the human skull. We demonstrate results with different parametrizations starting from one homogeneous skull conductivity value and ending with 8860 locally varying skull conductivity values. In addition to the skull conductivities, a homogeneous (bulk) conductivity for the scalp is solved in every test case.
- We show how the choice of the skull conductivity resolution (parametrization) in the EIT imaging affects the EEG source localization results. We input the EIT solved conductivities in the modelling of the EEG source imaging problem in order to reduce the source localization errors (that would otherwise occur due to the unknown conductivities). In the EEG source imaging, we used the dipole scan algorithm due to its simplicity and because it allowed direct quantication of the source localization error.
- This is, to the best of our knowledge, the first study that compares different skull conductivity resolutions (parametrizations) in the EIT head imaging, and subsequently quantifies the benefits of each EIT imaging result in terms of EEG source localization. The significance of our work is that it acts as the next step in defining a protocol for the preparation of patient-specific head conductivity models which are essential for accurate examination and systematic monitoring of brain activity with high-resolution EEG imaging techniques.

## 2 Theory

### 2.1 EIT observation model

The observation model of EIT can be written as

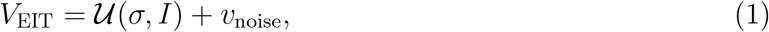

where *V*_EIT_ ∈ ℝ^*M*^ are the measured boundary voltages, 𝒰 (*σ, I*) is the mapping that connects the conductivity distribution *σ* and the pre-defined current injections *I* with the measured voltages, and *v*_noise_ represents random white measurement noise. For the numerical solution of the EIT problem, the conductivity distribution is discretized and approximated as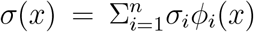, where *ϕ*_*i*_(*x*) can be linear basis functions, for example, and *n* is the total number of nodes of the finite element (FE) head model [44].

The mapping 𝒰 (*σ, I*) can be linearized with respect to the conductivity parameterization by using the Taylor series expansion

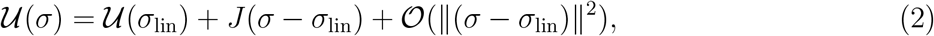

where *σ* ∈ ℝ^*n*^, *J* ∈ ℝ^*M×n*^, 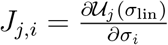 is the (*j, i*)th element of the Jacobian matrix [43] evaluated at a given linearization point *σ*_lin_ ∈ ℝ^*n*^.

From (1) and (2), we can approximate the observation model as

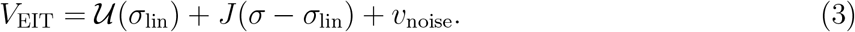

### 2.2 EIT inverse problem with box constraint

Here, the unknown conductivity parameters *σ* of (3) are solved with the help of a box constraint. The box constraint can be used to define the lower and upper bounds for the unknown conductivities (*σ*_*L*_ and *σ*_*U*_, respectively). The minimization problem and the corresponding minimization functional are of the form

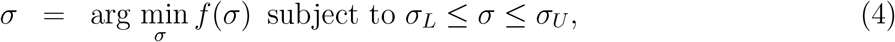

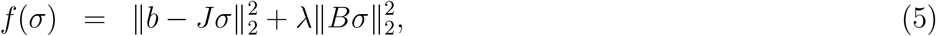

where *b* = *V*_EIT_ − 𝒰 (*σ*_lin_) + *Jσ*_lin_, *λ* is a regularization coefficient and *B* represents the conductivity (smoothness) prior. The problem is now solved iteratively by using algorithm 1.

#### Algorithm 1 Solve the EIT imaging problem

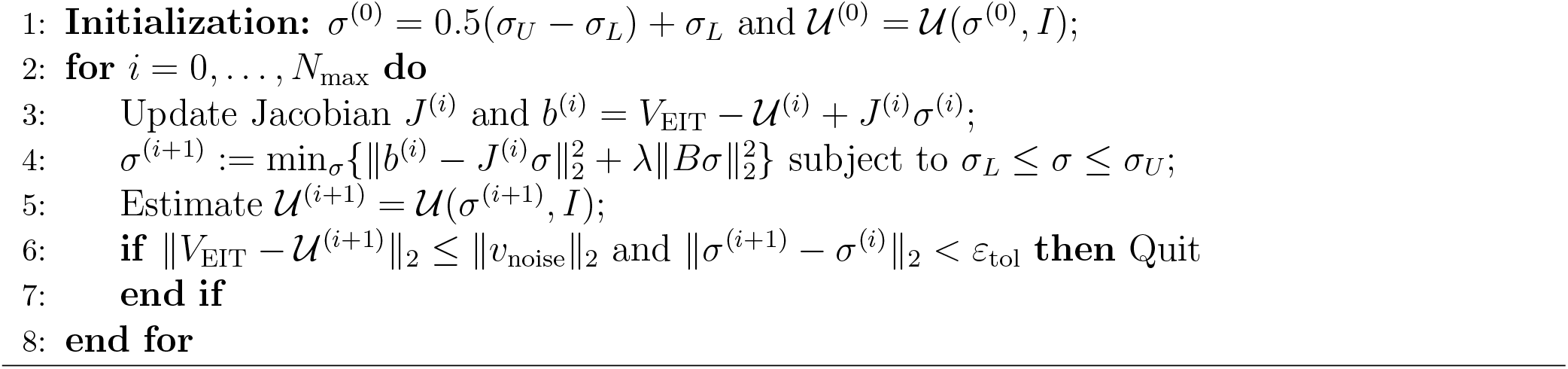

## 3 Methods

### 3.1 Numerical head models

A freely available numerical head model [41, 40] that consisted of scalp, skull, cerebro-spinal-fluid (CSF) and brain compartments was downloaded and then refined around the electrodes using a similar approach as in [39]. To simulate the EIT measurement data (forward problem), we used a mesh that consisted of 480471 elements joined in 85793 nodes. To solve the conductivities (inverse problem), we used a mesh with 168112 elements joined in 32331 nodes. For the measurements, 33 electrodes (of which the one in location FPZ was used for grounding) were placed on the scalp according to [15].

### 3.2 Simulation set-up in EIT forward problem

A locally varying (distributed) skull conductivity was created for testing the algorithm in reconstructing locally varying skull conductivity. This testing distribution was created based on [23] in which 20 bulk samples were drilled from a post-mortem skull and their conductivities were determined by using four electrode technique (please, see [23] for details). It was found that the conductivity values of the samples ranged from 0.0047 S/m (temporal bone) to 0.0735 S/m (occipital bone) [23]. Table 1 lists these conductivities and the scalp electrode locations under which the skull samples were taken. These locations are also shown in Fig. 1. For the simulation set-up (forward problem), these conductivity values were smoothened across the rest of skull to create the studied distributed skull conductivity, see Fig. 2(a).

**Table 1:**
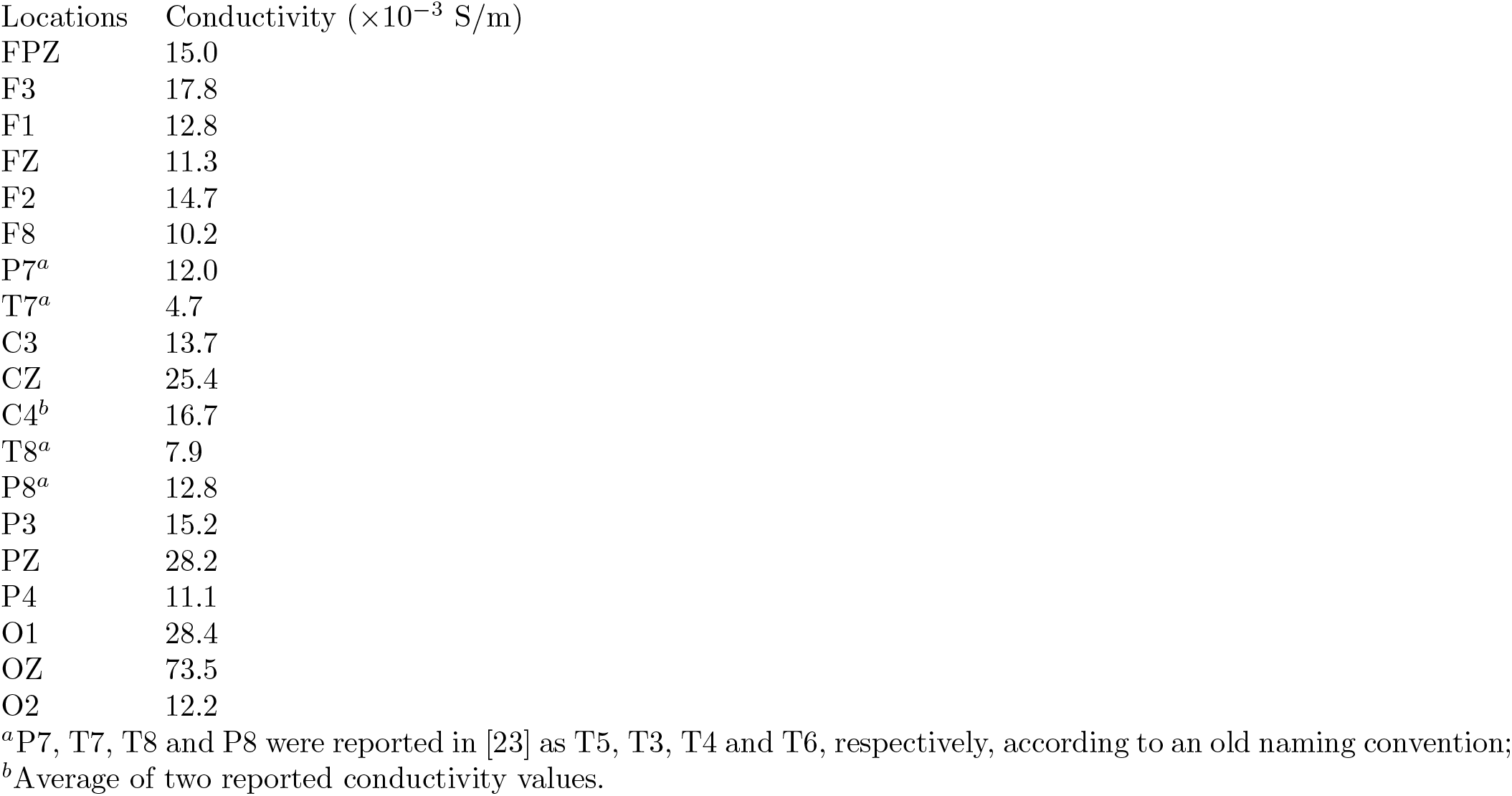
Electric conductivity values used in the test case for various locations of the skull. The values are based on [23].

**Figure 1:**
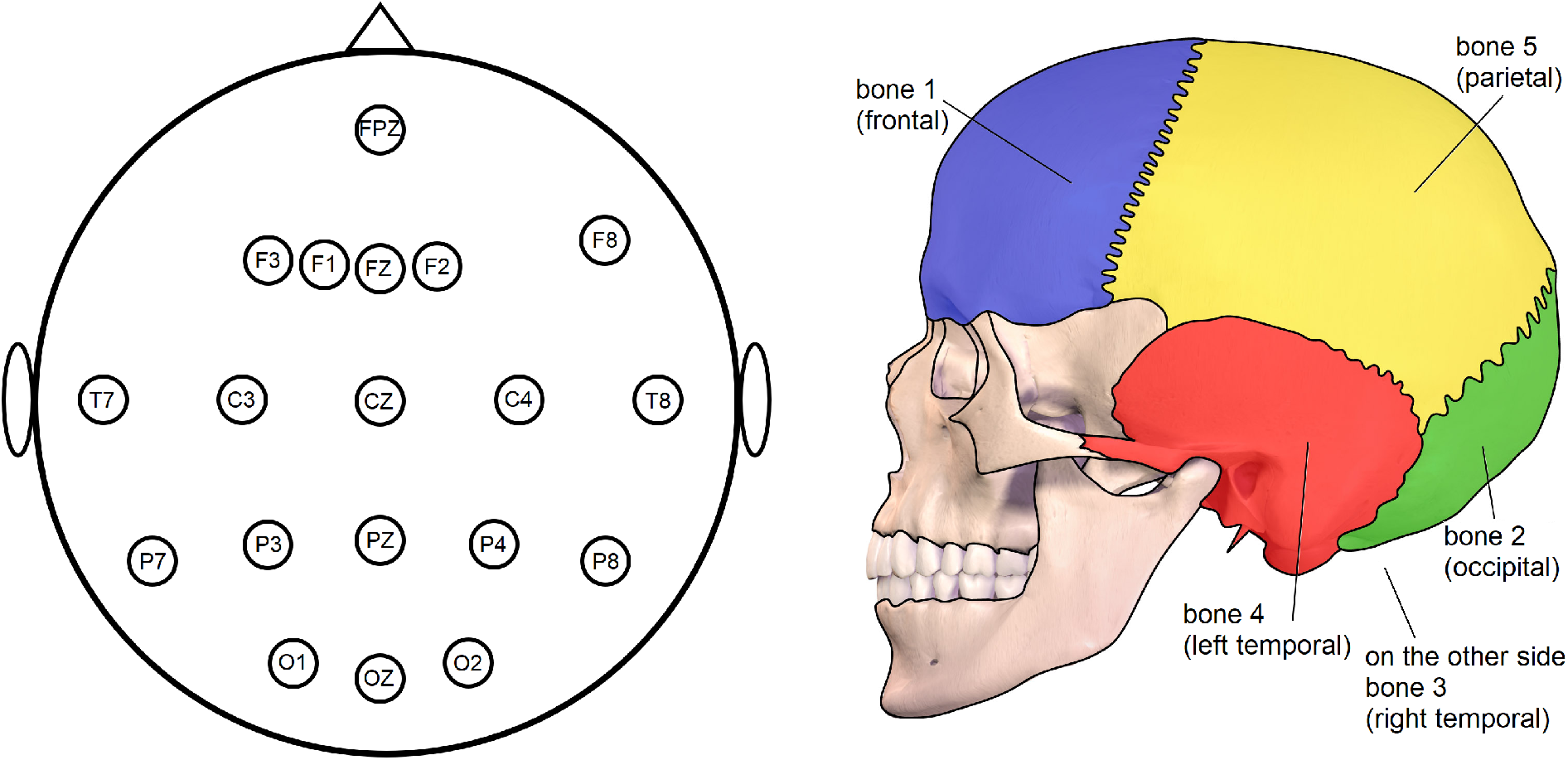
Left: This diagram shows the EEG electrode locations under which the skull samples were taken in [23]. Right: For the analysis in Section 4.3.2, the results were grouped based on the five main skull bones [1].

**Figure 2:**
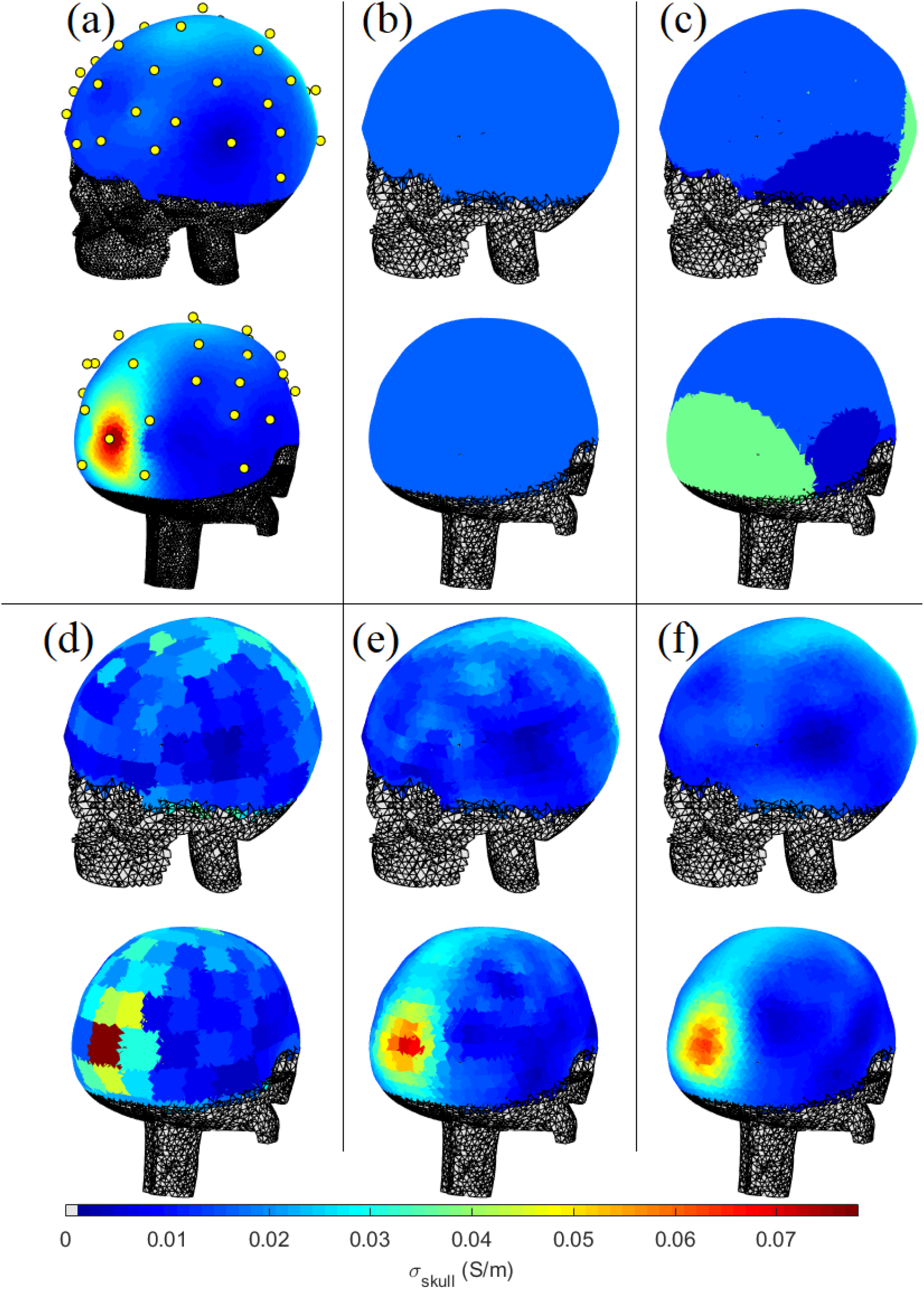
(a): True skull conductivity distribution used to simulate (forward) EIT data. Scalp electrodes are shown with yellow dots. (b): EIT solved (bulk) homogeneous skull conductivity. (c): EIT solved skull conductivities when one conductivity value is reconstructed for each of the 5 main bones of the head. (d)-(f): EIT imaging results when the skull conductivity parametrization is further refined to 172, 1064 and 8860 conductivity parameters, respectively.

For the rest of the tissues, we used the following (bulk) conductivity values (in S/m): *σ*_scalp_ = 0.43 [36], *σ*_CSF_ = 1.79 [6] and *σ*_brain_ = 0.33 [36].

The numerical EIT measurements were then carried out by injecting +1 mA and −1 mA currents through pairs of electrodes (one pair at a time) and measuring the resulting voltages against the ground electrode (FPZ electrode). Similarly as in [15], 31 current injections were used, and this resulted in a total of 992 voltage measurements of which 930 were subsequently used for the EIT analysis after exclusion of measurements on injection channels. Finally, 5% of simulated random white noise was added in the measurements.

### 3.3 Skull conductivity parametrization in EIT inverse problem

In this paper, we studied how different parametrizations affect the EIT inverse solution, and particularly how this solution improved EEG inverse source estimates when the EEG leadfield matrix (in the EEG inverse source problem) was constructed based on the EIT inverse solution. In order to study different parametrizations, we bundled nearby skull elements together and considered that every bundle had only one conductivity value. The computational benefit of bundling the elements in this way was the reduction of the number of unknowns in the EIT inverse problem.

In the first parametrization, the skull conductivity was considered simply as one (homogeneous) unknown parameter. In the second parametrization, the skull was divided anatomically into 5 pieces that approximately encompassed the frontal bone, left and right temporal bones, occipital bone and parietal bone, and one conductivity value was retrieved for each. The parametrization was further refined resulting in 172, 1064 and 8860 skull pieces (or number of unknowns). The lower and upper bounds of the box constraint were 0.001 S/m and 0.09 S/m, respectively.

In addition to skull, we reconstructed one homogeneous conductivity value for the scalp conductivity in each of the experiments. Since we were primaly interested in the skull conductivity (and different skull conductivity parametrizations), conductivities of CSF and brain were considered as known.

### 3.4 Jacobian matrix for the skull pieces

In the computations, the Jacobian matrix *J* ∈ ℝ^*M×n*^ (where *M* is the number of measurements and *n* total number of elements) needed to be re-defined for each of the skull parametrizations. First, the columns of the Jacobian matrix that corresponded to known conductivities were discarded reducing the size of the Jacobian matrix to *M × n*_sk_ where *n*_sk_ is the number of the elements in the skull compartment. For skull parametrization that had *c* unknown skull pieces, the columns of the Jacobian matrix that corresponded to a same piece were summed up, reducing the size of the *J* to *M × c*. In addition, the Jacobian has one column corresponding the scalp conductivity.

### 3.5 Normalized graph Laplacian

In our implementation, the matrix *B* was the normalized graph Laplacian [31] given by

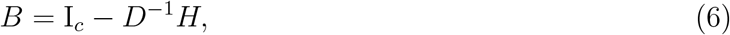

where I_*c*_ is the identity matrix of size *c, D* ∈ ℝ^*c×c*^ is a diagonal matrix with elements 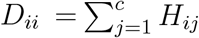, and matrix *H* ∈ ℝ^*c×c*^ has *H*_*ii*_ = 0 and non-zero elements for *i* ≠ *j*, 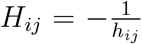, where *h*_*ij*_ is the distance between skull pieces *i* and *j* if these are connected, otherwise *H*_*ij*_ = 0. This choice of Laplacian ensures a smooth transition between neighboring pieces.

### 3.6 Implementation of box constraint

The problem (4) was solved by using the interior point method (IPM) [7]. For our implementation, the problem (4) was first reformulated as an unconstrained problem with the help of the logarithmic barrier method [7]

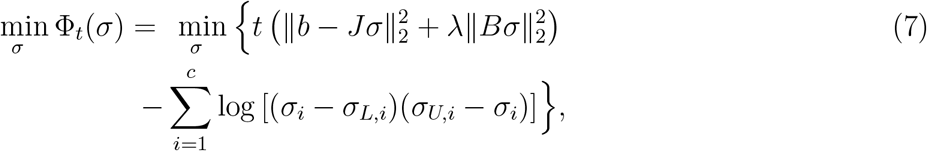

where *σ* ∈ ℝ^*c*^ is the unknown conductivity vector, *σ*_*L,i*_ and *σ*_*U,i*_ are the lower and upper conductivity limits in each segment.

The solution is then achieved by solving a sequence of unconstrained problems (7) for increasing values of *t >* 0. The IPM algorithm consists of an inner and outer loop [7]. In the inner loop, problem (7) is solved by keeping the value of *t >* 0 constant while in the outer loop, the value of *t* is updated. The IPM algorithm terminates based on a chosen stopping criterion. Here, we used the duality gap as in [20, 22].

#### 3.6.1 Stopping Criterion

As in [20], we use the difference between the values of functional of (eq. 4) and its dual problem, called duality gap, as a stopping criterion for the IMP algorithm. This stopping criterion relies on the fact that the Lagrange dual problem of problem (4) is always concave and yields lower bounds on the optimal value of convex problem (4) [7]. Here, we derive a Lagrange dual form of the convex problem (4) and we show how to estimate the duality gap.

To ease the derivation of the dual function, we re-write the cost function of problem (4) as

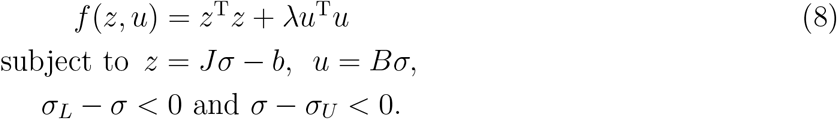

Then, the augmented Lagrangian is

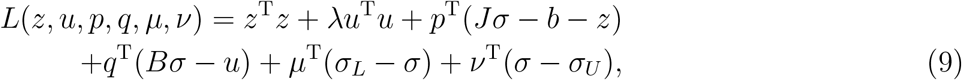

where *p, q, µ* ≥ 0, *ν* ≥ 0 are the Lagrange multipliers.

We estimate the Lagrange dual function,

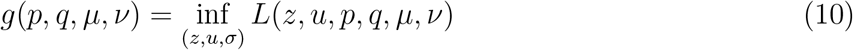

by minimizing the Lagrangian over *z, u* and *σ*. So, 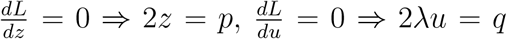 and 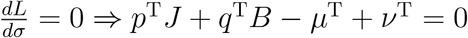 and the dual function becomes

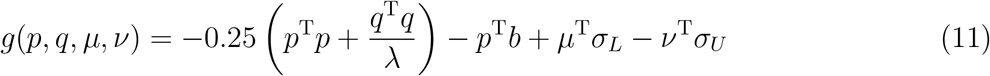

Therefore, the dual problem is

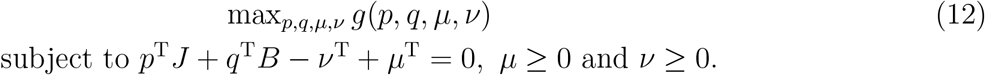

Duality gap is the difference between the value of *f* (*σ*) in equation (4) and its dual functional *g* given by

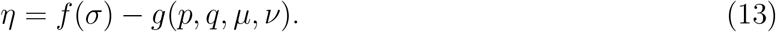

Given a solution *σ* for problem (4), we estimate *g* based on

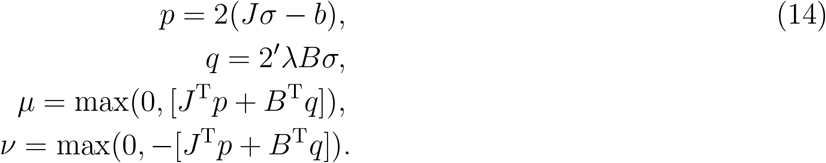

#### 3.6.2 Inner loop

Keeping *t* fixed, the problem (7) can be solved with the help of the Newton method [16] followed by backtracking line search (BLS) [7]. In particular, we estimated first the search direction defined as Δ*σ* ∈ ℝ^*c*^ by solving the linear system

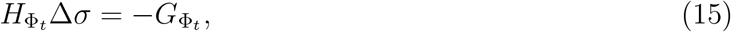

where 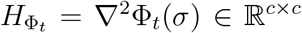 is the Hessian matrix and 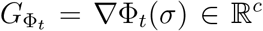 is the gradient of Φ_*t*_(*σ*), respectively. Then the updated solution is *σ* := *σ* + *s*Δ*σ* where *s >* 0 is the step size estimated by the BLS.

#### 3.6.3 Outer loop

The outer loop of the IPM updates the value of *t* [22, 20]. Here, we used

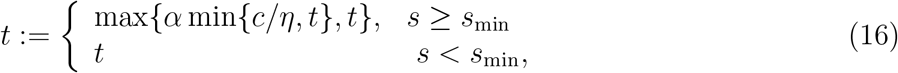

where *s* is the estimated step size and *s*_min_ = 0.7 is the threshold that determines whether *t* will be updated or not. This *t*-rule ensures that *t* stays constant until the cost function Φ_*t*_ (7) is nearly minimized, i.e. ∥∇Φ_*t*_∥_2_ ≃0.

## 4 Results and discussion

### Conductivity reconstructions from EIT data

The EIT reconstructions with the different skull conductivity parametrizations are shown in Fig. 2. Each of the results is an average of 5 solutions computed from EIT data with different noise realizations. Along with the presented skull conductivities, a homogeneous estimate for the scalp conductivity was solved. The true scalp conductivity value was 0.43 S/m whereas the reconstructed values were 0.3819 S/m, 0.3878 S/m, 0.4314 S/m, 0.4298 S/m and 0.4337 S/m for the results shown in Fig. 2 (b)-(f), respectively. It can be seen that the EIT solution resembles the more closely the true one the more there are skull conductivity parameters. Also the scalp conductivity estimate seems to be in general more accurate when the higher skull conductivity parametrizations are used. In the following, the notation *a+b parametrization* refers to *a* solved parameters for the scalp and *b* solved parameters for the skull.

*1+1 parametrization, Fig. 2(b):* In this parametrization one value was solved for both the scalp (0.3819 S/m) and skull (0.0159 S/m). The scalp conductivity is fairly far from the true one, and the skull conductivity seems to be almost an average of the values over the true distribution shown in Fig. 2 (and the measured values reported in Table 1). This result obviously is problematic around the very low conductivity areas (temporal bones) and the very high conductivity areas (in the occipital bone). Since a single conductivity value for the skull is not sufficient to explain the local conductivity variations in the skull, the reconstruction algorithm tries to tune also the scalp conductivity value to better match the EIT data. This result is an example of a case in which the parameterization is chosen poorly.

*1+5 parametrization, Fig. 2(c):* In this result the homogeneous scalp conductivty was 0.3878 S/m which is similar to the one in the previous parametrization. For the different bones, 0.0157 S/m was solved for frontal, 0.0377 S/m for occipital, 0.0062 S/m for right temporal, 0.0060 S/m for left temporal and 0.0157 S/m for parietal bone. As a rough sanity check these can be compared to the values reported in Table 1: the average of FPZ, FZ and F1-F3 conductivity samples 0.0143 S/m is close to the reconstructed frontal bone conductivity 0.0157 S/m; the average of O1, O2 and OZ samples 0.0380 S/m agrees with the reconstructed occipital bone conductivity 0.0377 S/m; the average of CZ, C3, C4, P3 and P4 samples 0.0164 S/m is close to the reconstructed parietal bone conductivity 0.0157 S/m; and the left/right temporal bone conductivities (T7 and T8) 0.0047 S/m and 0.0079 S/m are of the same order of magnitude as the reconstructed values 0.0062 S/m and 0.0060 S/m, respectively. Visually, one can say that the 1+5 parametrization works fairly well elsewhere except along the middle line of the skull and around the high conductivities in the occipital bone.

*1+172 parametrization, Fig 2(d):* In this parametrization, the skull was divided into 172 pieces (with average volume 2.2 cm^3^) each of which was reconstructed with a single conductivity value. In this case, the homogeneous scalp conductivity was correctly found (0.4314 S/m when the true value was 0.43 S/m). In addition, visually this parametrization seems to capture at least some of the local variations in the skull conductivity, e.g. the high conductivities in the occipital bone can be seen in one of the pieces. Obviously, this kind of coarse parametrization does not allow fine details to be captured.

*1+1064 parametrization, Fig. 2(d):* This parametrization was similar to the previous except now there were 1064 skull pieces (with average volume 0.37 cm^3^) for each of which a single conductivity value was solved. The solved scalp conductivity 0.4298 S/m was very close to the true one and also the reconstructed skull conductivity distribution resembles the original. However, one can still note that the high conductivity area at the occipital bone is smaller and the conductivity values are lower than in the true. Furthermore, the distribution of skull conductivities on top of the parietal bone is not quite right even though there is a significant improvement when compared to the other parametrizations.

*1+8860 parametrization, Fig. 2(e):* Here, the skull pieces were even smaller (with average volume 0.04 cm^3^) which enabled solving fine details in the skull conductivity profile. The scalp conductivity was found to be 0.4337 S/m which is slightly higher than the true one. The reconstructed skull conductivity follows very closely the true one, e.g. the local variations around the frontal bone were captured accurately. Again, only at the high conductive parts of the occipital bone there were minor errors: the high conductive area is slightly smaller than what it should be and the highest conductivity values are a bit too low. Nevertheless, the resemblance in general is excellent.

### 4.2 Discussion on EIT results

The numbers of unknowns in the different parametrizations were 2, 6, 173, 1065 and 8861 whereas the number of used EIT measurements was 930 in each of the case. Seemingly, to solve only 2 or 6 conductivity parameters one would not need such a number of measurements. However, the problem with using less data, say, one current injection with only few voltage measurements, is that the solution may be biased towards the local conductivity values close to the current injecting electrodes. For example, by reconstructing homogeneous scalp and skull conductivities with 30 EIT measurements from one current injection through electrodes F3 and FC1 results in 0.4472 S/m for scalp and 0.0155 S/m for skull conductivity. This skull conductivity value is now between the F1 and F3 samples (0.0128 S/m and 0.0178 S/m) given in Table 1. This is due to the fact that the sensitivity of the EIT measurements is the highest close to the current carrying electrodes [11]. Alternatively, if there are significant local conductivity variations around the current injecting electrodes, one can also get estimates that turn out to be misleading: when a similar experiment was conducted by having EIT data from only one current injection located around the left temporal bone (from T7 with sample value 0.0047 S/m to C3 with sample value 0.0137 S/m), a scalp conductivity of 0.2682 S/m and skull conductivity 0.0136 S/m were achieved. Here, the scalp conductivity value is significantly wrong (the algorithm is trying to use scalp conductivity value to explain the data), and skull conductivity (even though close to C3 value) is very far from T7 value. In order to not to be biased towards any local conductivity, it was deemed a good compromise to use all the available data. This also helped to make the EIT results comparable since all results were reconstructed from the same EIT data set.

For the 1+172, 1+1064 and 1+8860 parametrizations, a smoothness prior was needed in addition to the box constraint. Numerically, the imaging problem with the 1+172 parametrization is still over-determined and, thus, seemingly would not require regularization. However, it was found that the reconstructions with this parametrization were unstable with respect to noise and that regularization improved the EIT result. The prior was tailored for each case in such a way that the skull pieces, that were close to each other, were assumed to have smooth conductivity changes.

Based on these results, we can conclude that, by refining the skull conductivity parametrization, we can capture local conductivity variations in detail. Now, a question still remains: how much refinement there needs to be, i.e. how many skull conductivity parameters one needs to solve, in order to have also accurate EEG source imaging results? This question was investigated in the following section.

### 4.3 Source reconstructions from EEG data

#### 4.3.1 Different head conductivity models

To quantify the benefits of using the different skull conductivity parametrizations, we wanted to quantify the localization errors in the EEG source imaging when each of the parametrizations was used to construct the head volume conductor model in EEG source imaging problem. The EEG scalp recordings were first simulated (EEG forward problem) with the help of the lead field matrix ℒ (*σ*^acc^) that corresponded to the *accurate* skull conductivity (described in Section 3.2) and by placing single dipole sources (one at a time) in the brain, and finally the recordings were corrupted with 5 % simulated random white noise

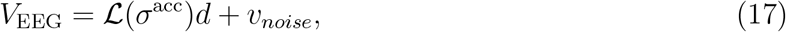

where 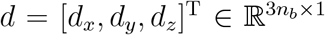 is the source configuration in the distributed source model, and *n*_*b*_ is the number of possible source locations in the distributed source model.

For the EEG source imaging, we constructed lead field matrices that corresponded the different EIT conductivity imaging results, i.e. ℒ (*σ*^(1+1)^), ℒ (*σ*^(1+5)^), ℒ (*σ*^(1+172)^), ℒ (*σ*^(1+1064)^) and ℒ (*σ*^(1+8860)^) corresponding to the results shown in Fig. 2. The superscripts refer to the corresponding conductivity parametrization. For the source reconstructions, we used the single dipole scan algorithm [30, 21] in which we fit a dipole for each source space node (considering that in every other node the dipoles are zero) and choose the dipole that achieves the smallest residual as the solution

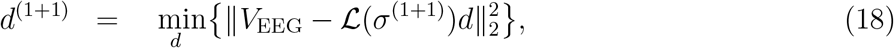

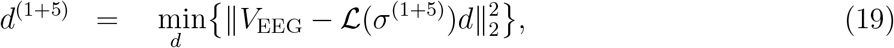

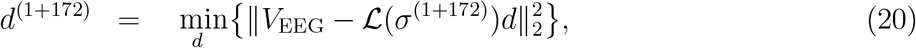

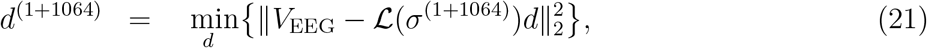

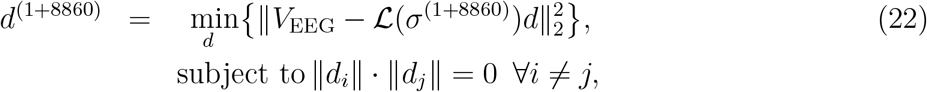

where ∥*d*_*i*_ ∥ is the strength of the dipole source at node *i*.

Since the standard methodology in EEG source imaging is to use various standardised bulk conductivity values for the tissues, we therefore also constructed such a standard model for comparison with bulk conductivity values obtained from literature. The idea here was to quantify the benefits of EIT resolved conductivity profiles against a standard one. In this *standard* head model, the bulk scalp conductivity was set 0.4137 S/m and bulk skull conductivity 0.016 S/m which were the ‘weighted mean values’ reported in [27]. Finally, we also used the *accurate* head model to reconstruct the sources as a reference.

#### 4.3.2 Source localization results

In this experiment, we placed one radial dipole source at a time in the brain that consisted of 5491 possible source locations, then computed the corresponding EEG data using (17), and finally reconstructed the sources by using the dipole scan algorithm once with each lead field model (18)-(22). Table 2 shows the average localization errors of the recontructed dipole sources with respect to their true locations. To ease the analysis, the results were categorized with the help of the locations of the true sources, namely, under which skull bone the source was located. In the notations, the numbers 1-5 correspond to frontal, occipital, right/left temporal and parietal bone, respectively (see, Fig. 1). Table 3 reports the percentages of cases in which the localization error was more than 6 mm.

**Table 2:**
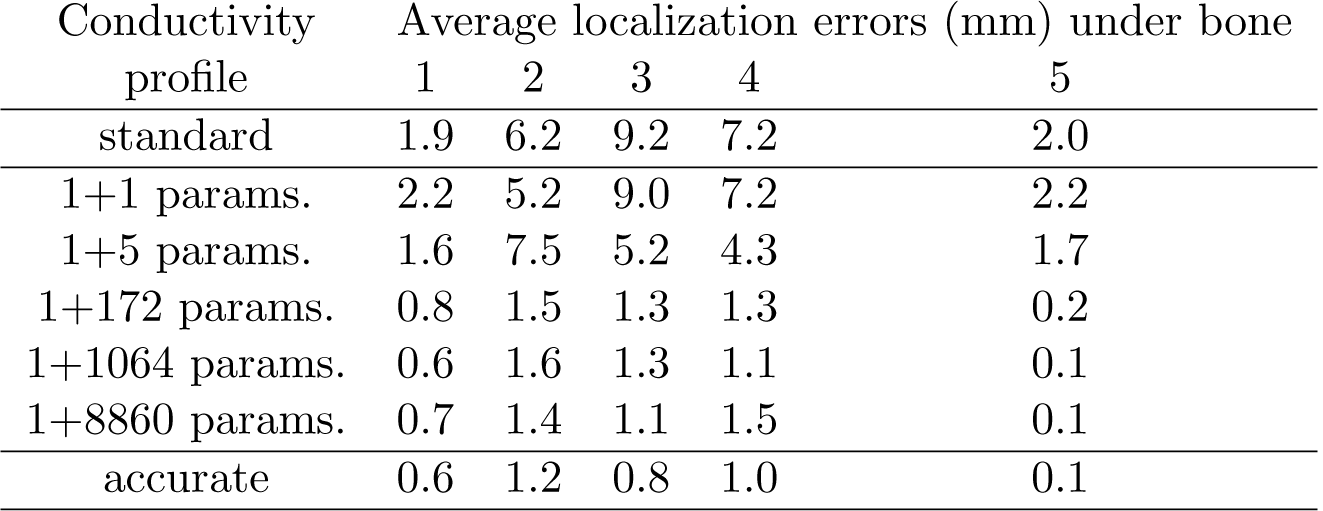
This table shows the averages of the localization errors when the different conductivity models were used in the EEG source imaging. The first column shows the number of parameters in the conductivity model, i.e. one homogeneous scalp conductivity and different numbers of skull conductivities. The localization errors (in millimetres) are given for sources that located under different skull bones: here 1 refers to frontal, 2 occipital, 3 right temporal, 4 left temporal and 5 to parietal bone.

| Conductivity<br>profile | Average localization errors (mm) under bone |  |  |  |  |
| --- | --- | --- | --- | --- | --- |
|  | 1 | 2 | 3 | 4 | 5 |
| standard | 1.9 | 6.2 | 9.2 | 7.2 | 2.0 |
| 1+1 params. | 2.2 | 5.2 | 9.0 | 7.2 | 2.2 |
| 1+5 params. | 1.6 | 7.5 | 5.2 | 4.3 | 1.7 |
| 1+172 params. | 0.8 | 1.5 | 1.3 | 1.3 | 0.2 |
| 1+1064 params. | 0.6 | 1.6 | 1.3 | 1.1 | 0.1 |
| 1+8860 params. | 0.7 | 1.4 | 1.1 | 1.5 | 0.1 |
| accurate | 0.6 | 1.2 | 0.8 | 1.0 | 0.1 |

**Table 3:** This table shows the percentages of cases in which the localization errors were above 6 mm when the different conductivity models were used in the EEG imaging. The first column shows the number of parameters in the conductivity model, i.e. one homogeneous scalp conductivity and different numbers of skull conductivities. The percentages are given for sources that located under different skull bones: here 1 refers to frontal, 2 occipital, 3 right temporal, 4 left temporal and 5 to parietal bone.

| Conductivity<br>profile | % of localization errors >6 mm under bone |  |  |  |  |
| --- | --- | --- | --- | --- | --- |
|  | 1 | 2 | 3 | 4 | 5 |
| standard | 16 | 50 | 84 | 71 | 16 |
| 1+1 params. | 19 | 42 | 85 | 71 | 18 |
| 1+5 params. | 13 | 65 | 47 | 40 | 13 |
| 1+172 params. | 7 | 10 | 8 | 9 | 1 |
| 1+1064 params. | 6 | 11 | 7 | 7 | 1 |
| 1+8860 params. | 6 | 9 | 6 | 10 | 1 |
| accurate | 5 | 7 | 5 | 7 | 1 |

### 4.4 Discussion on EEG results

Based on literature [35, 37], it is known that when a too high or a too low skull conductivity is used in the EEG source analysis the solutions will be biased. In particular, a too high skull conductivity pushes sources to locate deeper in the brain than they should. Analogously, a too low skull conductivity in EEG imaging brings sources too close to the skull. Furthermore, these localization errors are on average more prominent when a too high skull conductivity is used (rather than when a too low conductivity) [37].

From Tables 2 and 3, it can be seen that the standard model and the conductivity model based on the 1+1 parametrization had the worst performance of all models. In addition, both performed almost equally poorly. It is worth noting that the reference model was chosen to have a scalp conductivity (0.4137 S/m) that was very close to the true one (0.43 S/m) and skull conductivity (0.016 S/m) that was relatively close to many of the conductivity samples given in Table 1. In other words, without auxiliary information the scalp/skull conductivities could not be chosen as suitably as here. Based on these two models, one cannot recommend in general using a homogeneous (bulk) conductivity model for the skull.

Then, the EIT model based on the 1+5 parametrization worked mainly better than the two previously mentioned models. Particularly, improvements were observed in the region of the weakly conductive temporal bones. This parametrization worked poorer only when the sources were placed under the occipital bone. As was seen in Fig. 2, the 1+5 parametrization could not take properly into account the high local conductivities in this region which (in addition to the poor scalp conductivity estimate) resulted in the observed poor EEG performance.

By further refining the parametrization to 1+172, we see a particular improvement in localization errors in the occpital region. As was observed in Section 4.1, the 1+172 parametrization both provided an accurate scalp conductivity estimate and visually could represent the high occipital conductivities much better.

Then, even though Fig. 2 shows clear improvements in the resemblence of the skull conductivity distribution (when compared to the true) as the parametrization increases, these improvements do not seem to always propagate to the EEG results. Namely, the finest parametrizations 1+1064 and 1+8860, offer mainly marginal improvements to the previous. Therefore, one might argue that the 1+172 parametrization hits the sweet spot offering sufficient resolution for the EEG lead field model and not being computationally too demanding; it can be conditionally recommended as long as one takes its numerical sensitivity (see, Section 4.2) properly into account. One might also argue that if a higher number of electrodes or a more sophisticated EEG inverse algorithm with, say, an *-C*_1_ or a mixed norm prior were used, then the more refined parametrizations might show more benefits. The study of this hypothesis is out of the scope of this paper.

### 4.5 Our observations in brief

The main observation of our study are the following:

- In EIT scalp and skull conductivity imaging, as the parametrization was refined, we were able to retrieve better estimates for the bulk scalp conductivity and local variations in the skull conductivity.
- EIT conductivity imaging with appropriate parametrization can significantly improve the EEG lead field model and subsequently reduce the EEG source localization errors. In our example case, this was valid regardless in which part of the brain the sources located.
- A very commonly used conductivity parametrisation that has a bulk conductivity for both, scalp and skull, can result in very poor EEG localization errors. In our example case, the corresponding EEG localization errors were comparable to the standard model in which these conductivities were chosen based on literature. Therefore, this kind of bulk parametrization cannot be generally recommended.
- The brain sources that located below the parts of the skull with the lowest and the highest conductivities exhibited the highest localization errors in EEG imaging when either the standard or 1+1 model were used. Because of this, we can conclude, unsurprisingly, that more resolution in the EIT imaging is needed when there are significant local variations in the skull conductivity. In our example, this meant sources that located below the occipital and the temporal bones.
- The localization errors of brain sources reduced as the skull conductivity parametrization was refined, but this was observed only up to a point beyond which further refinement did not make much difference. In particular, 1+172 parametrization resulted in small EEG localization errors, and further refinement of the parametrization resulted in only marginal improvements in the EEG localization. This type of fairly coarse parametrization could therefore be recommended as long as its numerical stability is properly ensured with appropriate regularization techniques.

### 4.6 Discussion on further modelling of the skull

In reality, the skull consists of three layers: two layers of hard bone (with poor conductivity) and a layer of spongy bone (with higher conductivity) that is sandwitched in the between. In practice, however, the segmentatation of spongy bone is challenging, and many commonly used segmentation tools [32] do not have this option. Therefore, the necessity of the three-layer modelling of the skull has been investigated along with whole skull modelling approaches that can *effectively* model the skull as a single layer [10, 9, 29, 45]. In this paper, we introduced a whole skull modelling approach which allows local conductivity variations in the skull. As a result, the spongy bone areas do not need to be segmented, instead, the EIT solved locally varying conductivity takes the spongy bone regions into account with higher conductivity values.

## 5 Conclusions and future work

In this paper, we used a whole skull modelling that allowed local conductivity variations to be considered. Then, we solved using electrical impedance tomography with different parametrizations a locally varying skull conductivity profile and a bulk value for the scalp conductivity in a simulated human head. As was expected, the finer parametrizations were able to retrieve finer details of the skull conductivity. Furthermore, we quantified the benefit (in terms of source localization error) of inputting this information to a personalised lead field model in EEG source imaging.

The significance of our work is that it acts as the next step in defining a protocol for the preparation of patient-specific head conductivity models which are essential for accurate examination and systematic monitoring of brain activity with high-resolution EEG imaging techniques. Our future work consists of verifying the findings of this paper with dummy head models and laboratory experiments under different electrode settings and inverse algorithms.

## Acknowledgment

This research received funding from the ATTRACT project funded by the EC under Grant Agreement 777222, from the Academy of Finland post-doctoral program (project no. 316542), and from the Academy of Finland (project no. 336357, PROFI 6 - TAU Imaging Research Platform).

The authors wish to thank Dr. Nathan D. Smith for double checking the dual function.

## Footnotes

1 We wish to note that EIT can also be used as a stand-alone diagnostic tool [26, 33, 46, 2] to characterize strokes, for example. However, we do not aim to use EIT for diagnostics, thus the diagnostic papers are out of the scope of this work.

